# Inhibitory ultrapotent chemogenetics activate dopamine D1 receptor-expressing medium spiny neurons

**DOI:** 10.1101/2020.07.01.181925

**Authors:** Stephanie C. Gantz, Maria M. Ortiz, Andrew J. Belilos, Khaled Moussawi

## Abstract

Ultrapotent chemogenetics, including the chloride-permeable inhibitory PSAM^4^-GlyR receptor, were recently proposed as a powerful strategy to selectively control neuronal activity in awake, behaving animals. We aimed to validate the inhibitory function of PSAM^4^-GlyR in dopamine D1 receptor-expressing medium spiny neurons (D1-MSNs) in the ventral striatum. Activation of PSAM^4^-GlyR with the uPSEM^792^ ligand enhanced rather than suppressed the activity of D1-MSNs *in vivo* as indicated by increased c-fos expression in D1-MSNs. Whole-cell recordings in mouse brain slices showed that activation of PSAM^4^-GlyR did not inhibit firing of action potentials in D1-MSNs. Activation of PSAM^4^-GlyR depolarized D1-MSNs, attenuated GABAergic inhibition, and shifted the reversal potential of PSAM^4^-GlyR current to more depolarized potentials, perpetuating the depolarizing effect of receptor activation. The data show that ‘inhibitory’ PSAM^4^-GlyR chemogenetics may actually activate certain cell types, and highlight the pitfalls of utilizing chloride conductances to inhibit neurons.

## INTRODUCTION

Novel research tools like opto- and chemogenetics have been instrumental in dissecting brain circuits and understanding their relevance to normal and maladaptive behaviors. However, like any new technology, these tools have inherent limitations that could confound data interpretation.

The leading chemogenetic approach, Designer Receptors Exclusively Activated by Designer Drugs (DREADDs), acts through engineered G protein-coupled receptors, which are selectively activated by clozapine N-oxide (CNO) (1). However, DREADDs may disrupt signaling from endogenous G protein-coupled receptors when expressed at high levels (2), and CNO has been found to be metabolized peripherally to clozapine, which binds to many receptors in the brain, potentially resulting in off-target effects (3, 4). A more recent chemogenetic approach, Pharmacologically Selective Actuator/Effector Module (PSAM/PSEM), acts through engineered ligand-gated ion channels that are activated by PSEM agonists (5, 6). The inhibitory PSAM^4^-GlyR is a chimeric protein of a modified α7 nicotinic acetylcholine receptor ligand binding domain fused to the chloride-permeable glycine receptor ion pore domain (GlyR). After intracranial injection and expression of a virus encoding PSAM^4^-GlyR, application of ultrapotent agonist (e.g., uPSEM^792^) activates PSAM^4^-GlyR, which in principle selectively inhibits PSAM^4^-GlyR-expressing neurons (6).

In this study, we aimed to validate the inhibitory function of PSAM^4^-GlyR in dopamine D1 receptor-expressing medium spiny neurons (D1-MSNs) in the ventral striatum. Intraperitoneal (i.p.) injection of uPSEM^792^ enhanced the expression of the immediate early gene c-fos in PSAM^4^-GlyR-expressing D1-MSNs, indicative of *in vivo* activation rather than inhibition of these neurons. Using whole-cell recordings in acute brain slices, we found that activation of PSAM^4^-GlyR with uPSEM^792^ decreased membrane resistance, induced an inward current in voltage-clamp, and depolarized the membrane potential in current-clamp, sometimes resulting in depolarization block. The majority of neurons maintained action potential firing to somatic current injection. Further, we found that chloride influx via PSAM^4^-GlyR activation shifted its reversal potential to more positive values further exacerbating the depolarizing effect, and attenuated GABAergic inhibition of D1-MSNs.

## RESULTS

### Selective expression of PSAM^4^-GlyR in D1-MSNs

To first validate the selective expression of PSAM^4^-GlyR in ventral striatum D1-MSNs, prodynorphin-Cre mice were crossed with a Cre-reporter (Ai9) mouse line, resulting in labeling of neurons containing Cre-recombinase (i.e., D1-MSNs) with the fluorescent reporter tdTomato (7-9). These mice received bilateral stereotaxic microinjections of AAV-syn-flex-PSAM^4^-GlyR-IRES-eGFP. PSAM^4^-GlyR expression was restricted to tdTomato^+^ neurons which confirmed selective Cre-dependent expression of PSAM^4^-GlyR in D1-MSNs (Figure 1A).

**Figure 1.**
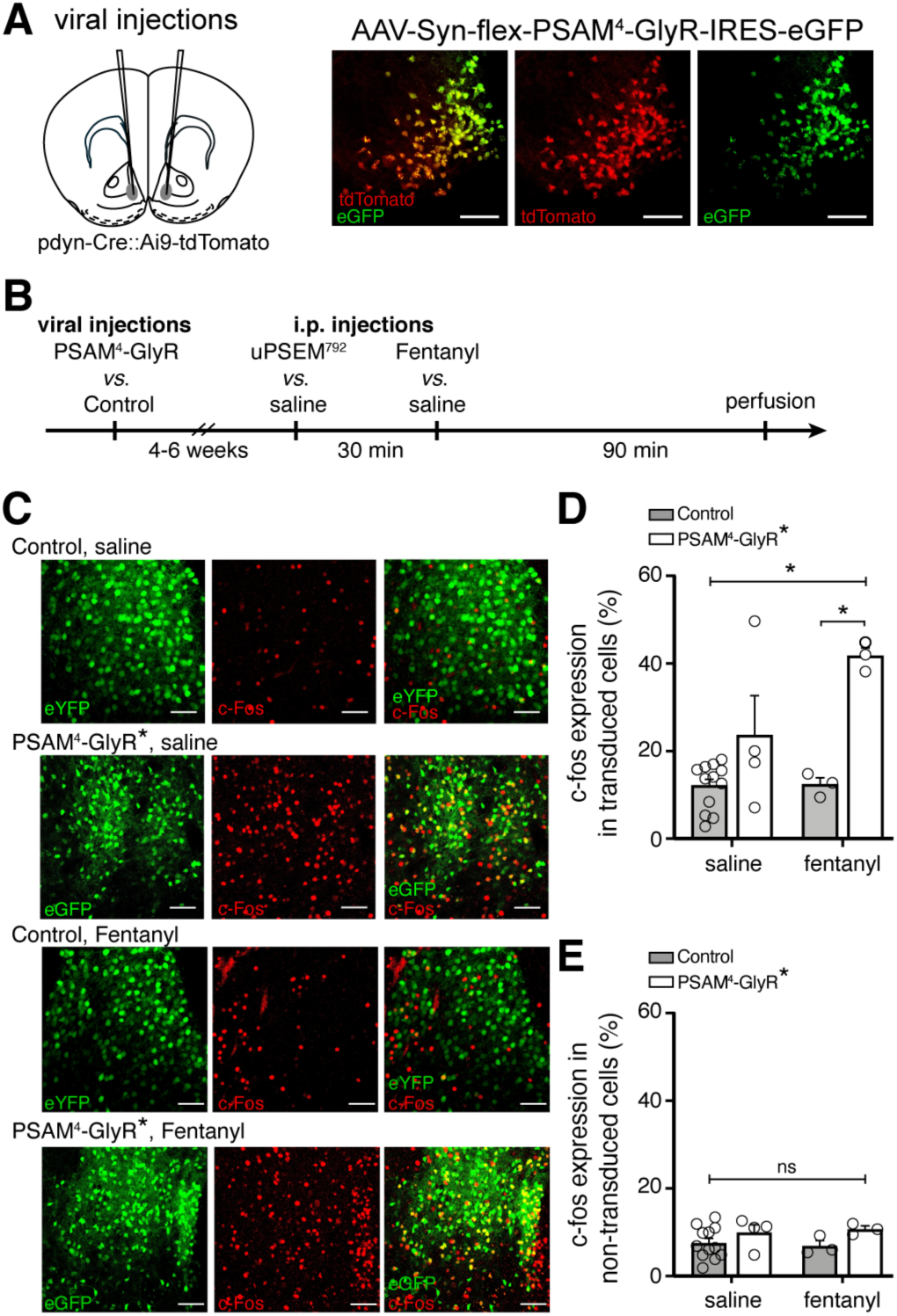
*In vivo* activation of PSAM^4^-GlyR enhances c-fos expression in D1-MSNs. (**A**) Left, cartoon of viral injection site in the ventral striatum. Right, representative confocal images of Cre-dependent PSAM^4^-GlyR expression (eGFP) in Cre-expressing D1-MSNs (tdTomato). (**B**) Experimental design for measuring c-fos expression after activation of PSAM^4^-GlyR. **(C)** Representative maximum intensity Z-stack confocal images of c-fos immunostaining (red) in D1-MSNs with or without activation of PSAM^4^-GlyR, after saline *vs*. fentanyl injection (0.2 mg/kg, i.p.). PSAM^4^-GlyR* refers to PSAM^4^-GlyR activated with uPSEM^792^ (3 mg/kg, i.p.). The ‘Control’ group includes pooled data from mice with (control virus + uPSEM^792^ 1^st^ i.p. injection) and (PSAM^4^-GlyR virus + saline 1^st^ i.p. injection). Scale bars: 50 µm. (**D**) Activation of PSAM^4^-GlyR with uPSEM^792^ increased c-fos expression in PSAM^4^-GlyR expressing D1-MSNs compared to control (two-way ANOVA with Sidak’s multiple comparisons test). Data are presented as % colocalization of c-fos and eGFP/eYFP^+^ in transduced D1-MSNs. (**E**) Activation of PSAM^4^-GlyR showed a trend towards a significant increase in c-fos expression in non-transduced cells (calculated as the number of c-fos expressing non-transduced cells / total number of non-transduced cells; two-way ANOVA, PSAM^4^-GlyR activation effect: *p* = 0.07). Data represent mean ± SEM. * indicates statistical significance, ns denotes not significant.

### PSAM^4^-GlyR enhances rather than suppresses c-fos expression in D1-MSNs *in vivo*

The immediate early gene c-fos is commonly used as a surrogate marker for neuronal activation (10-13), and opioid exposure has been shown to increase the expression of c-fos and other immediate early genes in striatal neurons including D1-MSNs (14-17). Given the proposed role of PSAM^4^-GlyR activation in silencing neurons (5, 6), we initially hypothesized that activating PSAM^4^-GlyR *in vivo* will suppress D1-MSNs activity, and therefore reduce both baseline c-fos as well as opioid-induced increase in c-fos expression. Prodynorphin-Cre mice received bilateral injections of either AAV-syn-flex-PSAM^4^-GlyR-IRES-eGFP (PSAM^4^-GlyR) (*n* = 12) or AAV-EF1α-DIO-eYFP (*n* = 10). Then, after 4-6 weeks, mice were injected with uPSEM^792^ (i.p., 3 mg/kg) or saline, followed 30 minutes later by fentanyl (i.p., 0.2 mg/kg) or saline (Figure 1B). Mice were perfused and their brains collected for c-fos immunostaining 90 min later (120 min after the first i.p. injection). Transduced neurons were identified by expression of the fluorescent reporter (eGFP or eYFP, green, Figure 1C). Contrary to expectation, activation of PSAM^4^-GlyR with i.p. uPSEM^792^ increased c-fos expression in D1-MSNs expressing PSAM^4^-GlyR (Figure 1D). Two-way ANOVA showed a significant effect of PSAM^4^-GlyR activation on c-fos expression (F_1, 18_ = 22.23, *p* = 0.0002), a significant effect of fentanyl injection (F_1, 18_ = 4.51, *p* = 0.048), and a significant PSAM^4^-GlyR activation × fentanyl interaction (F_1, 18_ = 4.23, *p* = 0.05). The experimental groups included 6-7 mice/group, but some mice were eliminated because of failure of viral expression. While the resulting *n* in some groups was low, it was sufficient to reject the initial hypothesis that PSAM^4^-GlyR activation reduces c-fos expression in D1-MSNs. There was no difference in c-fos expression in non-transduced cells (two-way ANOVA; PSAM^4^-GlyR activation effect: F_1, 18_ = 3.68, *p* = 0.07; Figure 1E). These results show that activation of PSAM^4^-GlyR increased c-fos expression in transduced cells and therefore activated rather than inhibited D1-MSNs *in vivo*.

### PSAM^4^-GlyR activation depolarizes D1-MSNs

To test the effects of PSAM^4^-GlyR on electrophysiological properties of D1-MSNs, PSAM^4^-GlyR-encoding virus was injected into the ventral striatum of prodynorphin-Cre mice. After 4-6 weeks, whole-cell patch-clamp recordings (V_hold_ -88 mV) were made from eGFP^+^ neurons in acute mouse brain slices. PSAM^4^-GlyR was activated by application of the ligand uPSEM^792^. To control for any off-target effects of uPSEM^792^, recordings were also made from control neurons (neighboring eGFP^-^ neurons and eYFP^+^ neurons from mice that received bilateral injections of AAV-EF1α-DIO-eYFP). In control neurons, 10 or 50 nM uPSEM^792^ had no effect on basal whole-cell current (10 nM: -2.8 ± 7.7 pA, *n* = 6, *p* = 0.73, paired *t*-test; 50 nM: -9.5 ± 9.4 pA, *n* = 14, *p* = 0.24; Figure 2A). In PSAM^4^-GlyR^+^ neurons, 10 nM uPSEM^792^ produced an inward current (−43.0 ± 10.1 pA, *n* = 21, *p* = 0.0004, paired *t*-test; Figure 2A and B), consistent with the original report on PSAM^4^-GlyRs (6). Higher concentrations of uPSEM^792^ produced substantially larger inward currents (50 nM: -275.5 ± 57.1 pA, *n* = 18, *p* = 0.0002, paired *t*-test; 100 nM: -410.1 ± 137.8 pA, *n* = 6, *p* = 0.031; Figure 2A and B). To assay membrane resistance, the instantaneous change in current in response to a 10 mV step (V_hold_ -88 to -78 mV) was measured before and after uPSEM^792^ application (Figure 2C). The basal membrane resistance of D1-MSNs was 78.6 ± 5.5 MΩ, consistent with a previous report using similar recording conditions (18), and was reduced by activation of PSAM^4^-GlyR (10 nM: 6.5 ± 3.3% decrease, *n* = 9, *p* = 0.08, paired *t*-test; 50 nM: 29.6 ± 5.3% decrease, *n* = 18, *p*< 0.0001; Figure 2D).

**Figure 2.**
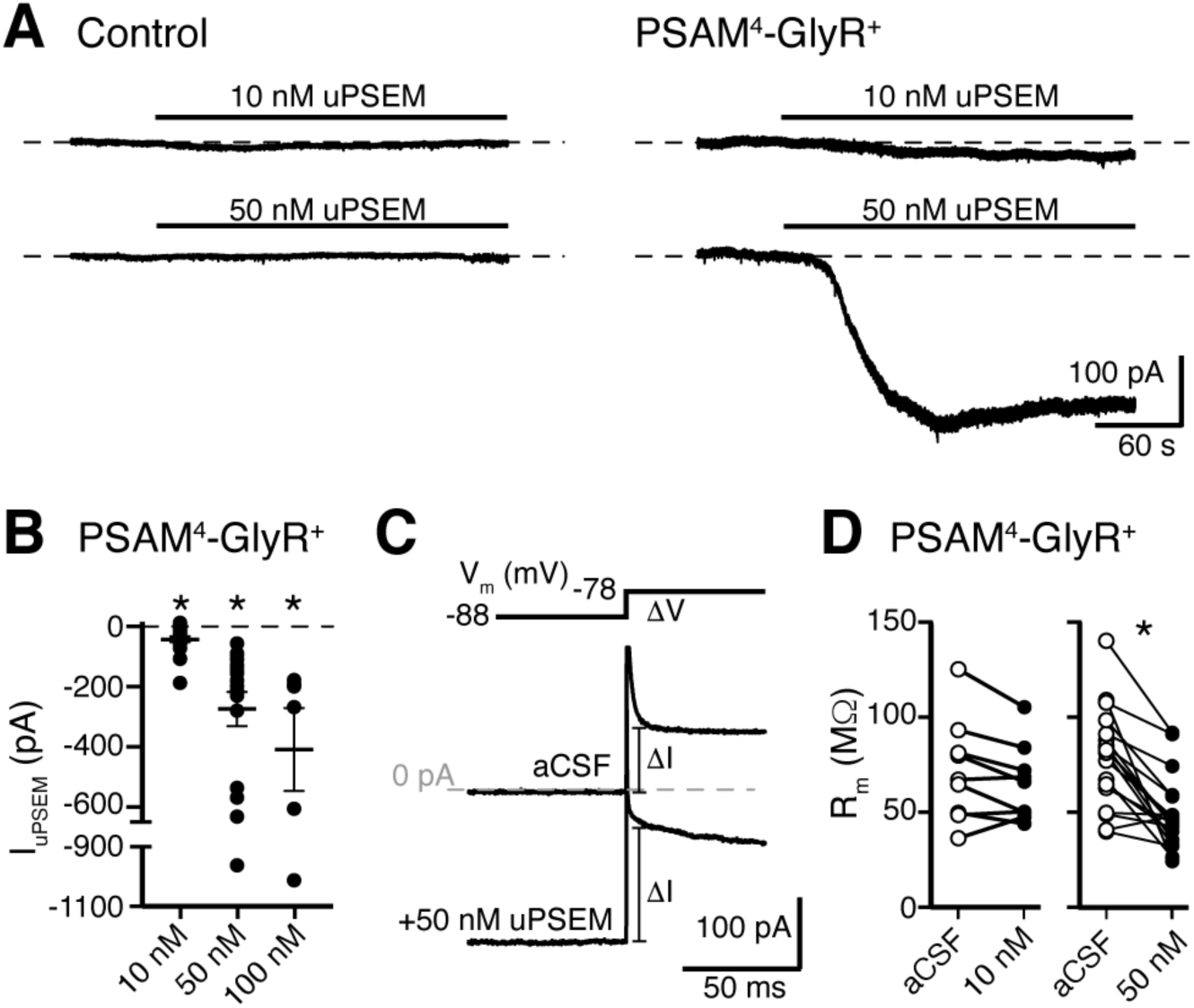
Activation of PSAM^4^-GlyR in D1-MSNs produces an inward current. (**A**) Representative traces of whole-cell voltage-clamp recordings (V_hold_ -88 mV) demonstrate no effect of uPSEM^792^ (10 or 50 nM) in control neurons. In PSAM^4^-GlyR^+^ neurons, 10 or 50 nM uPSEM^792^ produced an inward current. Dashed line is baseline whole-cell current for ease of visualization. (**B**) Plot of the magnitude of the inward current produced by 10, 50, or 100 nM uPSEM^792^ in PSAM^4^-GlyR^+^ neurons. Line and error bars represent means ± SEM. (**C**) To measure membrane resistance (R_m_), a 10 mV voltage step (−88 to - 78 mV) was made in aCSF and during the uPSEM^792^-induced inward current, and the instantaneous change in current following the capacitive transient was measured (ΔI). Representative traces are shown below the voltage step command. Dashed line is 0 pA. (**D**) In PSAM^4^-GlyR^+^ neurons, 50 nM uPSEM^792^ significantly decreased R_m_ (paired *t*-test, *p*< 0.0001) while 10 nM showed a trend towards lower R_m_ (paired *t*-test, *p* = 0.08). * indicates statistical significance, ns denotes not significant.

### PSAM^4^-GlyR does not silence D1-MSN action potential firing

We next determined the effect of PSAM^4^-GlyR activation on neuronal excitability. Action potential firing was evoked by somatic current injections (50 pA, 2 s) during whole-cell current-clamp recordings from control and PSAM^4^-GlyR^+^ neurons. PSAM^4^-GlyR activation by uPSEM^792^ significantly depolarized the membrane potential of PSAM^4^-GlyR^+^ neurons compared to control neurons (Figure 3A and B). Two-way ANOVA showed a significant effect of PSAM^4^-GlyR activation on membrane potential (F_1, 37_ = 9.70, *p* = 0.004), and a significant effect of uPSEM^792^ concentration (F_1, 37_ = 4.96, *p* = 0.03). Relative to the resting membrane potential (−83.3 ± 0.7 mV), the depolarization induced by 10 and 50 nM uPSEM^792^ was 6.4 ± 2.0 mV (*n* = 14, *p* = 0.007, paired *t*-test) and 20.9 ± 3.4 mV (*n* = 16, *p*< 0.0001), respectively.

**Figure 3.**
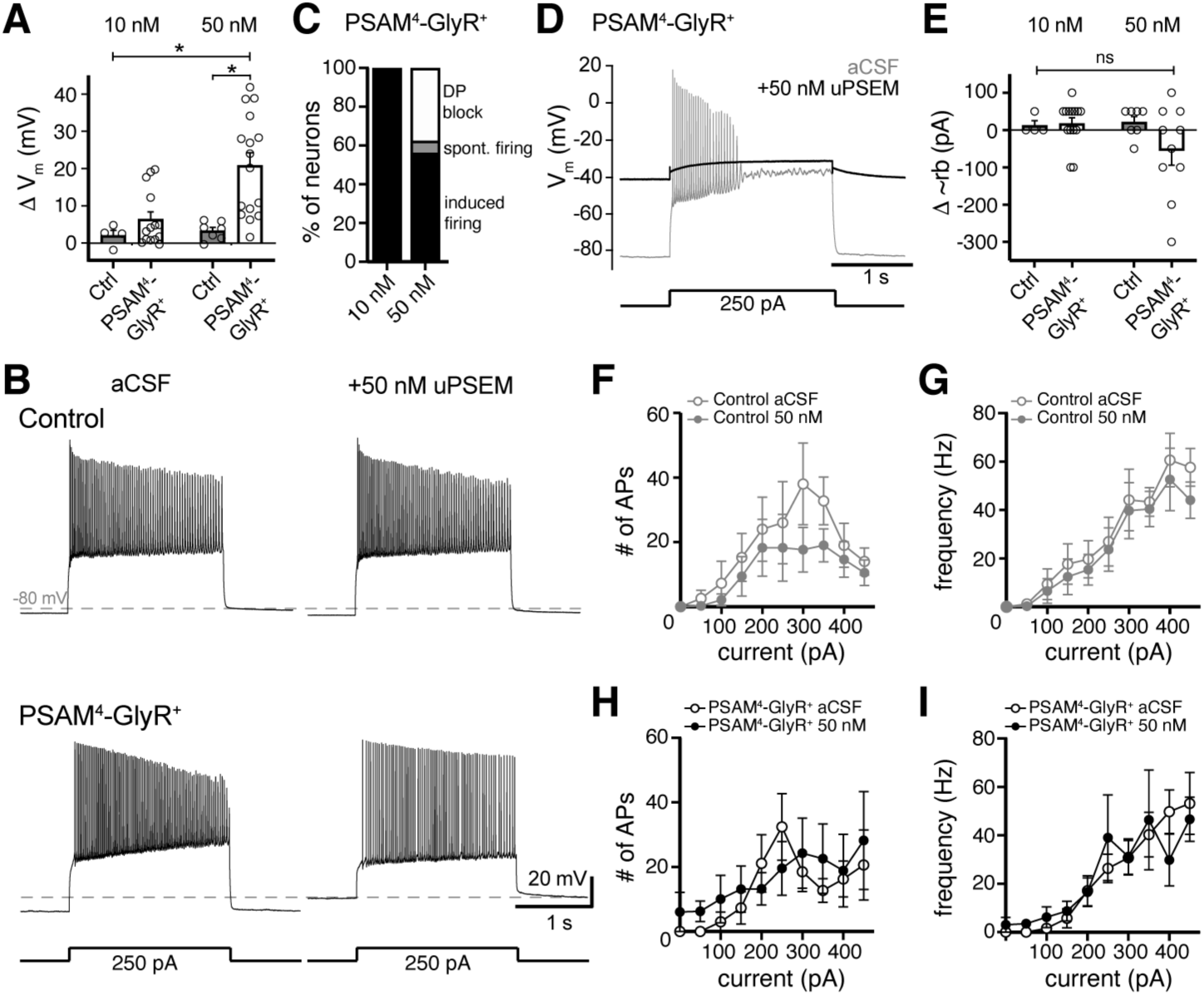
Activation of PSAM^4^-GlyR depolarizes D1-MSNs and does not inhibit firing. (**A**) In PSAM^4^-GlyR^+^ neurons, activation of PSAM^4^-GlyR with uPSEM^792^ (10, 50 nM) significantly depolarized the membrane potential (V_m_) compared to control (non-PSAM^4^-GlyR) neurons, measured in current-clamp (two-way ANOVA with Sidak’s multiple comparisons test). (**B**) Representative traces of current-clamp recordings from a control (top) or PSAM^4^-GlyR^+^ neuron (bottom). Action potential firing was evoked by somatic current injection (250 pA, 2 s) in aCSF (left) and 50 nM uPSEM^792^ (right). Dashed line is -80 mV. (**C**) Distribution of firing response of PSAM^4^-GlyR^+^ neurons after 10 or 50 nM uPSEM^792^ application. In 10 nM uPSEM^792^, 14/14 neurons continued to fire with current injection. In 50 nM, 9/16 neurons fired with current injection (induced firing), 1/16 fired spontaneously without a current injection (spont. firing), and 6/16 went into depolarization block (DP block). (**D**) Representative traces of current-clamp recordings from a PSAM^4^-GlyR^+^ neuron that went into depolarization block with uPSEM^792^. Response to somatic current injection (250 pA, 2 s) in aCSF (grey) and after uPSEM^792^ application (black). The membrane potential in uPSEM^792^ was more depolarized than the potential where the neuron entered depolarization block in aCSF. Dashed line is -80 mV. (**E**) Plot of change in the minimum current needed to induce firing (approximate rheobase: ∼rb) induced by uPSEM^792^ (10, 50 nM) in control (grey) and PSAM^4^-GlyR^+^ (black). No significant change of ∼rb was observed. (**F**) Plot of number of action potentials (APs) fired versus injected current in control neurons in aCSF (open) and in uPSEM^792^ (closed). No significant difference was observed. (**G**) Plot of AP firing frequency versus injected current in control neurons in aCSF (open) and in uPSEM^792^ (closed). No significant difference was observed. (**H**) Plot of number of APs fired versus injected current in PSAM^4^-GlyR^+^ neurons in aCSF (open) and in uPSEM^792^ (closed). No significant difference was observed. (**I**) Plot of AP firing frequency versus injected current in PSAM^4^-GlyR^+^ neurons in aCSF (open) and in uPSEM^792^ (closed). No significant difference was observed. Line and error bars represent means ± SEM, * indicates statistical significance, ns denotes not significant.

In control neurons, uPSEM^792^ (50 nM) did not change the number of evoked action potentials (*n* = 4-7, ANOVA mixed-effects analysis; no significant group effect: F_1, 6_ = 0.83, *p* = 0.40, and no interaction: F_2.1, 8.73_ = 1.30, *p* = 0.32; Figure 3B and F), or firing frequency (no significant group effect: F_1, 6_ = 0.29, *p* = 0.61, and no interaction: F_2.97, 12.55_ = 2.78, *p* = 0.08; Figure 3G). In PSAM^4^-GlyR^+^ neurons, all neurons treated with 10 nM uPSEM^792^ continued to fire action potentials with somatic current injections (*n* = 14, Figure 3C). In 50 nM uPSEM^792^, 10 of 16 neurons continued to fire action potentials with current injection and 6 of 16 neurons were depolarized sufficiently to enter into depolarization block upon uPSEM^792^ application (Figure 3C and D). In the cells that continued to fire, there was no change in the minimum positive current needed to evoke firing (approximation of rheobase) upon activation of PSAM^4^-GlyR (two-way ANOVA; PSAM^4^-GlyR activation effect: F _1, 31_ = 1.17, *p* = 0.29; uPSEM^792^ concentration effect: F _1, 31_ =1.45, *p* = 0.24; Figure 3E). Moreover, activation of PSAM^4^-GlyR with 50 nM UPSEM^792^ did not change the number of action potentials (*n* = 5-10, ANOVA mixed-effects analysis; no significant group effect: F_1, 9_ = 0.20, *p* = 0.66, and no interaction: F_9, 49_ = 0.96, *p* = 0.48; Figure 3H) or firing frequency (no significant group effect: F_1, 9_ = 0.05, *p* = 0.82, and no interaction: F_9, 48_ = 0.66, *p* = 0.48; Figure 3I). These results demonstrate that activation of PSAM^4^-GlyR depolarized the membrane potential and that the magnitude of the decrease in membrane resistance produced by opening of PSAM^4^-GlyR channels was not sufficient to inhibit neuronal firing via electrical shunting.

### Sustained activation of PSAM^4^-GlyR results in runaway depolarization of D1-MSNs

Depolarization from the resting membrane potential (−83 mV) by activation of PSAM^4^-GlyR was expected given the calculated reversal potential of the channel (−62 mV, see Methods). However, in most D1-MSNs (∼57%), the depolarization overshot the expected reversal potential to potentials more positive than -62 mV (Figure 3A). Therefore, we examined the current-voltage relationship of PSAM^4^-GlyR. Voltage steps (from V_hold_ -88 mV, 1 s, -118 to -28 mV, 10 mV increments) were made in aCSF and uPSEM^792^, and PSAM^4^-GlyR current was isolated by subtraction (Figure 4A). In control neurons, uPSEM^792^ did not produce substantial current across the voltage range (Figure 4A and C). In PSAM^4^-GlyR^+^ neurons, uPSEM^792^ produced a transient current with the onset of the voltage step (inset “1”, Figure 4A) that relaxed towards a steady-state current during the voltage step (inset “2”, Figure 4A). The transient current reversed polarity at -61.4 mV (*n* = 21, CI: -64.0 to -59.0 mV, linear regression, r^2^ = 0.58, F_1, 103_ = 142.5, *p*< 0.0001), consistent with the calculated reversal potential of the PSAM^4^-GlyR channel (see Methods). Transient PSAM^4^-GlyR current was inward at potentials more negative than -61 mV, and outward at potentials less negative than -61 mV (Figure 4C). At steady-state, the reversal potential of PSAM^4^-GlyR current showed a significant rightward shift to -44.2 mV (*n* = 21, CI: -48.3 to -38.2 mV, linear regression, r^2^ = 0.39, F_1, 103_ = 66.9, *p*< 0.0001; paired *t*-test, *t* = 5.73, *p*< 0.0001, Figure 4D and E). Further, the steady-state current showed a U-shaped current-voltage relationship (Figure 4D), and the outward current component was significantly smaller than the transient outward current (49.8 ± 38.4 pA *vs*. 307.1 ± 62.0 pA at V_hold_ -28 mV respectively, unpaired *t*-test, *p*< 0.0001; Figure 4D). Thus, in D1-MSNs, the majority of outward current carried by PSAM^4^-GlyR channels was transient. During the firing cycle, membrane hyperpolarization via PSAM^4^-GlyR activation is expected to be minimal or transient, and thus, PSAM^4^-GlyR activation is predominantly depolarizing.

**Figure 4.**
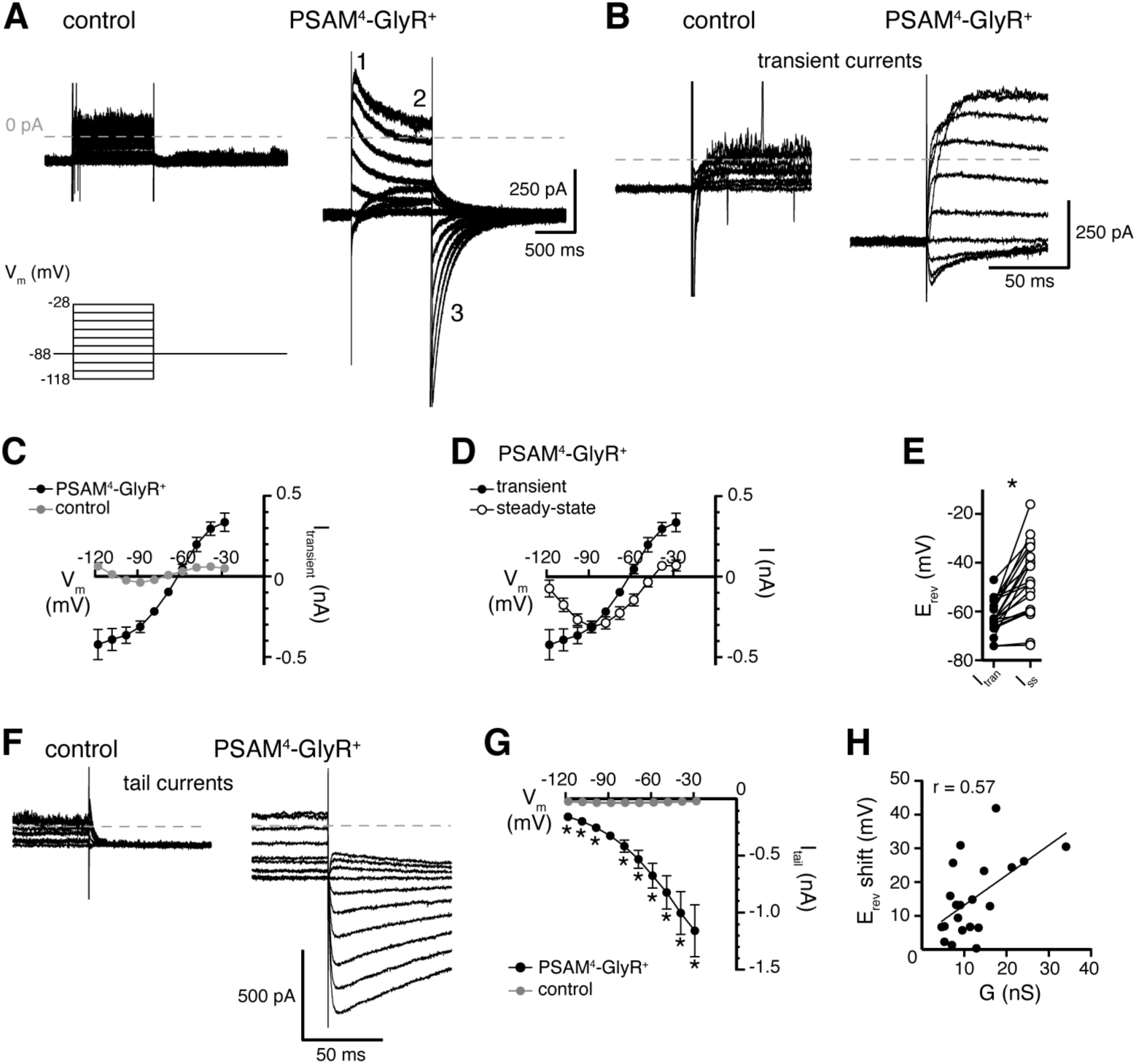
Current-voltage relationship of PSAM^4^-GlyR current shows rightward shift in reversal potential and transient or small outward currents at depolarized potentials. (**A**) Current was recorded in response to voltage steps (1 s, -118 to -28 mV, 10 mV) from V_hold_ -88 mV in aCSF and after application of uPSEM^792^ (50 or 100 nM). uPSEM^792^-induced current was isolated by subtraction. Representative traces of uPSEM^792^-induced current during voltage steps in a control (left) and a PSAM^4^-GlyR^+^ neuron (right). Inset numbers by trace indicate transient (1), steady-state (2) and tail (3) currents. (**B**) Expanded timescale of transient currents (inset “1” in A) in a control and a PSAM^4^-GlyR^+^ neuron. (**C**) Plot of current-voltage relationship of the transient current (I_transient_) in control (*n* = 11) *vs*. PSAM^4^-GlyR^+^ neurons (*n* = 21). The reversal potential of transient current in PSAM^4^-GlyR^+^ neurons was -61 mV. (**D**) Plot of current-voltage relationship of the transient (inset “1” in A, same as in C) *vs*. steady-state uPSEM^792^-induced (inset “2” in A) currents in PSAM^4^-GlyR^+^ neurons. At steady state, the reversal potential of uPSEM^792^-induced current shifted to -44 mV. (**E**) Plot of the pairwise shift of reversal potential (E_rev_) between transient (I_tran_) and steady-state currents (I_ss_). (**F**) Expanded timescale of tail currents (inset “3” in A) in a control and a PSAM^4^-GlyR^+^ neuron. (**G**) Plot of the amplitude of tail currents measured at V_hold_ -88 mV (I_tail_) versus holding potential of the preceding voltage step in control and PSAM^4^-GlyR^+^ neurons. uPSEM^792^-induced tail current at -88 mV demonstrated robust augmentation with prior depolarization and reduction with prior hyperpolarization. (**H**) The magnitude of the rightward shift in reversal potential between transient and steady-state current was positively correlated with the PSAM^4^-GlyR conductance (G in nS). Line and error bars represent means ± SEM, * indicates statistical significance.

### Instability of chloride equilibrium underlies PSAM^4^-GlyR-mediated depolarization

In order to gain a mechanistic understanding of PSAM^4^-GlyR-mediated depolarization, we examined tail currents generated by stepping back to a holding potential of -88 mV from preceding voltage steps. In principle, relaxation of outward currents during voltage steps could reflect fewer available channels (e.g., channel desensitization, inactivation, or voltage-dependent block), which would be expected to result in smaller tail currents. Alternatively, the relaxation of outward currents could be a manifestation of reduced driving force of chloride influx through PSAM^4^-GlyR channels due to the accumulation of intracellular chloride over the course of the voltage steps, which would be expected to result in larger tail currents. Indeed, many reports show that sustained activation of chloride conductances including GABA_A_ receptors (19-21), glycine receptors (22), and Halorhodopsin (23, 24) can lead to chloride influx that overwhelms homeostatic mechanisms to pump chloride out, thereby elevating the intracellular chloride concentration. An increase in intracellular chloride manifests as a rightward shift of the chloride reversal potential, commensurate with our observation of a rightward shift of the PSAM^4^-GlyR reversal potential from -61 mV to -44 mV. Further, the magnitude of the tail current increased substantially with preceding depolarizing steps (Figure 4F and G; RM two-way ANOVA, significant PSAM^4^-GlyR activation effect: F_1, 30_ = 18.05, *p* = 0.0002; significant PSAM^4^-GlyR activation × voltage step interaction: F_9, 270_ = 9.55, *p*< 0.0001; Dunnett’s multiple comparisons test shows that tail currents from every preceding voltage step were significantly different from current at -88 mV without a preceding voltage step). These data reveal that the decline in outward current was not due to fewer available channels, but was consistent with a change in the driving force and shift in the reversal potential during the depolarizing voltage steps. In addition, the magnitude of the shift in reversal potential was positively correlated with the conductance of PSAM^4^-GlyR, measured at V_hold_ -88 mV (Pearson’s correlation, r = 0.57, *p* = 0.0008, *n* = 21; Figure 4H). The tail current analysis also suggested some intrinsic voltage-sensitivity of PSAM^4^-GlyR. The magnitude of tail currents decreased with preceding hyperpolarizing steps (e.g., -98, -108, or -118 mV) and increased with preceding depolarizing steps (e.g., -78 or -68 mV; Figure 4F and G), suggesting that hyperpolarization reduced the number of available channels, or that depolarization increased the number of available channels. Taken together, the results suggest that activation of PSAM^4^-GlyR in D1-MSNs is largely depolarizing at sub- and suprathreshold membrane potentials likely due to two mechanisms: increased channel availability with depolarization due to intrinsic voltage-sensitive mechanisms, and more prominently, a rightward shift of PSAM^4^-GlyR reversal potential due to accumulation of intracellular chloride at depolarized potentials.

### Loss of GABAergic inhibition following activation of PSAM^4^-GlyR

Shifts in intracellular chloride concentration have been shown to result in “apparent cross-desensitization” of glycine- and GABA-A receptors whereby sustained activation of one chloride-permeable channel (e.g., glycine channel) results in reduced currents carried by another chloride-permeable channel (e.g., GABA-A channel) and vice versa (22). Therefore, we examined GABA-A receptor-mediated synaptic currents in PSAM^4^-GlyR^+^ neurons following activation of PSAM^4^-GlyR. A bipolar stimulating electrode was placed in the brain slice and GABA-A receptor-mediated synaptic currents were electrically evoked. The same voltage steps were made (as to assess the current-voltage relationship of PSAM^4^-GlyR current) and GABA-A receptor-mediated synaptic currents were evoked 850 ms into the voltage step where uPSEM^792^ current reached steady-state (single electrical pulse, 0.1 ms, Figure 5A and B). In neurons that showed a rightward shift in reversal potential of PSAM^4^-GlyR (> 5 mV shift, *n* = 5, E_rev_ shift = 17.3 ± 6.3 mV), the amplitude of the GABA-A receptor-mediated synaptic currents significantly decreased upon activation of PSAM^4^-GlyR compared with aCSF (−33.4 ± 6.0 pA *vs*. -7.8 ± 1.3 pA at V_hold_ -88 mV, paired *t*-test: *p* = 0.01; Figure 5C). Further, the reversal potential of GABA-A receptor-mediated synaptic currents showed a significant rightward shift from -60.9 mV in aCSF to -51.6 mV upon activation of PSAM^4^-GlyR (aCSF: CI: -64.8 to -57.9 mV, linear regression, r^2^ = 0.73, F_1, 13_ = 34.5, *p* < 0.0001; uPSEM^792^: CI: -57.7 to -42.7 mV, r^2^ = 0.66, F_1, 5_ = 9.6, *p* = 0.027; test of equal intercepts: F _1, 19_ = 13.3, *p* = 0.002; Figure 5C). In neurons that did not show a shift in reversal potential of PSAM^4^-GlyR (*n* = 3, E_rev_ shift = 2.1 ± 1.0 mV), there was no change in the reversal potential or amplitude of GABA-A receptor-mediated synaptic currents (aCSF: GABA-A E_rev_ = -64.7 mV, CI: - 68.5 to -62.1 mV, linear regression, r^2^ = 0.91, F_1, 7_ = 68.8, *p* < 0.0001; uPSEM^792^: GABA-A E_rev_ = -64.9 mV, CI: -66.7 to -63.4 mV, r^2^ = 0.96, F_1, 4_ = 87.5, *p* = 0.0007; Figure 5D). These results indicate that activation of PSAM^4^-GlyR reduced GABAergic inhibition of D1-MSNs.

**Figure 5.**
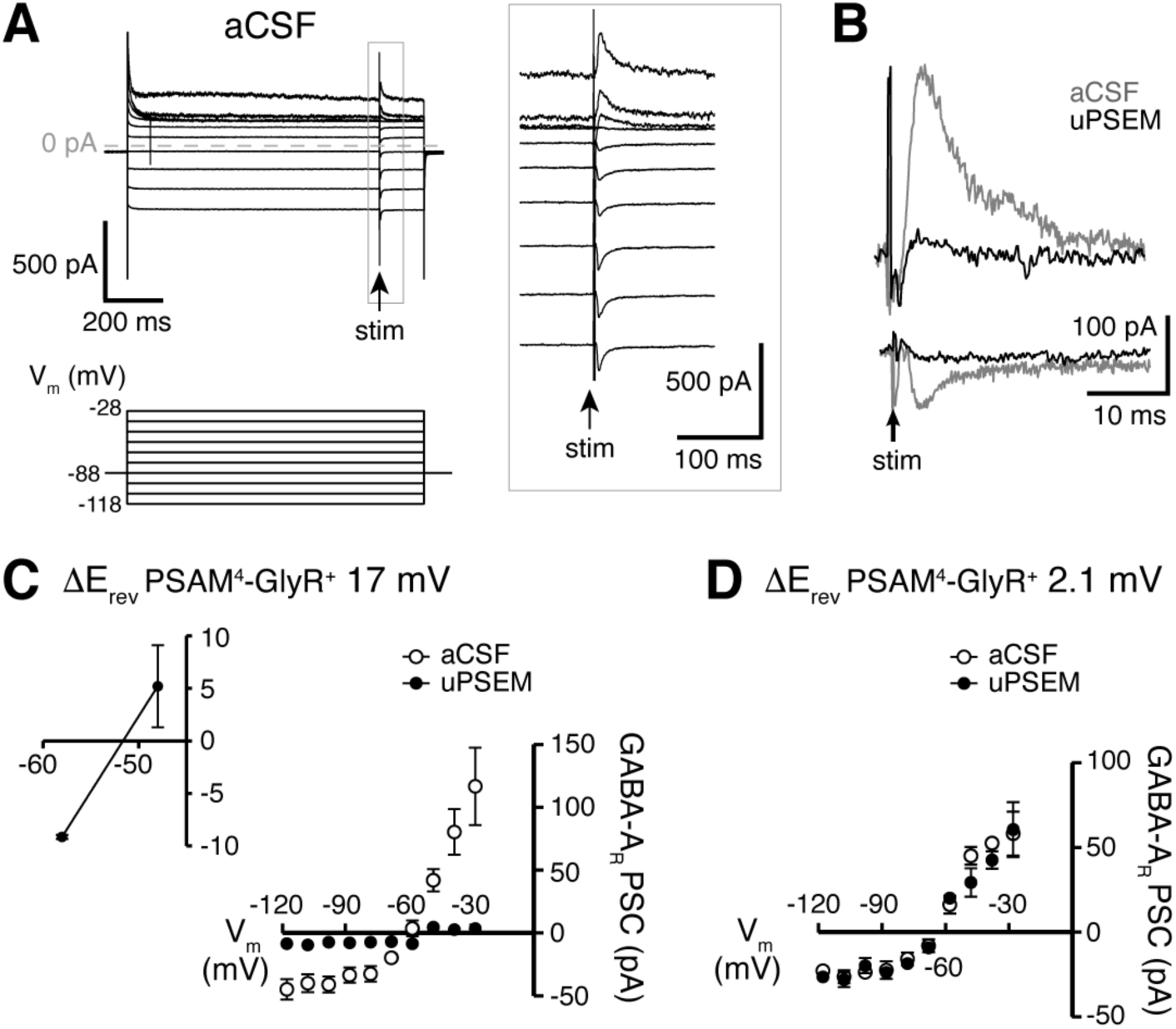
PSAM^4^-GlyR activation in D1-MSNs reduces GABAergic synaptic inhibition. (**A**) Representative traces of GABA-A receptor-mediated synaptic currents in a PSAM^4^-GlyR^+^ neuron in aCSF, evoked during voltage steps from V_hold_ -88 mV (voltage steps: 1 s, -118 to -28 mV, 10 mV; electrical stimulation at 850 ms into voltage steps). Panel on the right shows the traces in expanded timescale corresponding to the grey rectangle on the left. (**B**) Representative traces of GABA-A receptor-mediated currents evoked in the same neuron from V_hold_ -28 mV (outward) and -118 mV (inward) in aCSF (grey) and uPSEM^792^ (black). **(C)** Plot of the current-voltage relationship of the GABA-A receptor synaptic currents in aCSF and uPSEM^792^ in neurons where the reversal potential of PSAM^4^-GlyR current shifted by > 5 mV (*n* = 5, average shift of 17.0 mV). The reversal potential of GABA-A receptor-mediated current was -60.9 mV in aCSF and -51.6 mV in uPSEM^792^. Inset (top left) shows values of GABA-A receptor synaptic currents in uPSEM^792^ near the reversal potential in an expanded scale. (**D**) Plot of the current-voltage relationship of the GABA-A receptor-mediated synaptic current in aCSF and uPSEM^792^ in neurons where the reversal potential of PSAM^4^-GlyR current did not shift (*n* = 3, average shift of 2.1 mV). The reversal potential of GABA-A receptor-mediated synaptic currents was similar in aCSF (−64.7 mV) and uPSEM^792^ (−64.9 mV). Line and error bars represent means ± SEM, * indicates statistical significance.

## DISCUSSION

Strategies to manipulate neuronal activity in defined cell populations have become instrumental in mapping neural circuits and correlating neuronal and circuit activity with behavior. Engineered receptors to promote excitation (e.g., Channelrhodopsin-2, G_q_-DREADDs) have been largely successful, albeit with careful consideration of ligand-related off-target effects (3, 4). In contrast, silencing of neuronal firing has been uniquely challenging, especially when relying on chloride conductances (25). The utility of a chloride conductance like PSAM^4^-GlyR to silence firing relies on hyperpolarization of the membrane at potentials more positive than the chloride reversal potential and/or the efficacy of the channels to shunt membrane depolarization (26).

The present study shows that PSAM^4^-GlyR in D1-MSNs is predominantly depolarizing, with limited capacity to hyperpolarize these neurons even at more depolarized potentials. When activated near the resting membrane potential (−83 mV), PSAM^4^-GlyR passes inward current and depolarizes D1-MSNs. When activated at potentials more depolarized than the chloride reversal potential, PSAM^4^-GlyR passes transient or small outward currents, likely due to influx and accumulation of intracellular chloride that overwhelms endogenous mechanisms to restore the chloride gradient (19-24).

More commonly, chloride conductances inhibit action potential firing through electrical shunting. Opening channels decreases membrane resistance which reduces (shunts) depolarization in response to inward current. While there was a significant decrease in membrane resistance associated with opening PSAM^4^-GlyR in D1-MSNs (∼30%), it was not sufficient to prevent suprathreshold depolarization by current injection. The majority of neurons were equally capable of firing action potentials to current injection. Despite using high viral titer in our experiments, the decrease in membrane resistance we observed was lower than previously reported in cortical layer 2/3 neurons where PSAM^4^-GlyR activation suppressed firing (6). While increasing PSAM^4^-GlyR conductance by further increasing expression levels could theoretically improve shunting efficacy, the data in this study show a strong correlation between PSAM^4^-GlyR conductance and the rightward shift in PSAM^4^-GlyR reversal potential, suggesting that increasing expression levels will only promote further depolarization with PSAM^4^-GlyR activation.

*In vivo*, D1-MSNs oscillate between higher probability, less excitable ‘down-states’ (∼-77 mV) and lower probability, more excitable brief depolarization-plateaus referred to as ‘up-states’ (−50 to -70 mV, 100-1000 ms), which facilitate action potential firing (27, 28). Our current-clamp data show that activation of PSAM^4^-GlyR depolarizes D1-MSNs by ∼20 mV. This implies that *in vivo*, PSAM^4^-GlyR activation will effectively increase the probability and duration of ‘up-states’, and hence increase the probability of D1-MSN activation. In addition, the results show that PSAM^4^-GlyR activation results in ‘apparent cross-desensitization’ and profound reduction of inhibitory GABA-A synaptic currents onto D1-MSNs. Given the critical role of local GABAergic inhibition in the striatum in regulating neuronal activity (29, 30), PSAM^4^-GlyR triggered reduction of GABAergic inhibition will further increase the likelihood of D1-MSNs activation rather than inhibition. Thus, increased c-fos expression in D1-MSNs following *in vivo* activation of PSAM^4^-GlyR is likely caused by PSAM^4^-GlyR-induced depolarization and loss of GABAergic inhibition, and reflects increased neuronal activity of D1-MSNs *in vivo*.

Finally, PSAM^4^-GlyR-induced depolarization may allow robust calcium influx via voltage-gated calcium channels and NMDA receptors relieved from voltage-dependent magnesium pore-block (31, 32). Calcium influx activates calcium/calmodulin-dependent protein kinases (CaMKs), calcium response elements in genes, and various other signaling cascades (33, 34), which are known to influence synaptic plasticity and behavior (35, 36). Therefore, it is possible that when used to silence neurons, PSAM^4^-GlyR activation could potentially confound the interpretation of experimental results due to depolarization-induced calcium influx, independent of PSAM^4^-GlyR’s effect on action potential firing like in the case of depolarization-block.

In summary, activation of PSAM^4^-GlyR expressed in D1-MSNs in the ventral striatum enhanced neuronal activity through direct depolarization and did not suppress action potential firing via membrane shunting. The results of our study show that the PSAM^4^-GlyR approach to silence neurons may not be suitable for all cell types, and highlight the need to validate the inhibition of neuronal firing by PSAM^4^-GlyR in the cell type of interest prior to behavioral studies. More broadly, these data demonstrate that achieving neuronal silencing with chloride conductances continues to be challenging, and may result in unexpected neuronal activation.

## MATERIALS AND METHODS

### Subjects

All experimental procedures were conducted in accordance with the guidelines of the National Institutes of Health Guide for the Care and use of Laboratory Animals, and approved by the Animal Care and Use Committee of the National Institute on Drug Abuse. We used adult female and male prodynorphin-Cre mice (pdyn-Cre; 8-12 weeks old, breeding facility at the Intramural Research Program, National Institute on Drug Abuse) for electrophysiology and c-fos experiments. We crossed pdyn-Cre and Ai9 Rosa-tdTomato mice to validate selectivity of viral transduction (Figure 1A). Mice were group housed four per cage and maintained under a 12-h light cycle at 21±2 °C. Food and water were freely available.

### Stereotaxic Virus Injection

AAV-SYN-flex-PSAM^4^-GlyR-IRES-EGFP was a gift from Scott Sternson (Addgene viral prep # 119741-AAV5; http://n2t.net/addgene:119741; RRID:Addgene_119741; >1×10^13^ vg/ml). Control virus was AAV1-EF1α-DIO-eYFP (>1×10^13^ vg/ml, University of North Carolina Viral Core, NC). Mice were anesthetized with an i.p. injection of a cocktail of ketamine (100 mg/kg) and xylazine (10 mg/kg) and secured in a stereotaxic frame (David Kopf Instruments, CA). PSAM^4^-GlyR and control viruses were injected (0.2-0.5 µl, 0.05 µl/min) into the medial ventral striatum (targeted coordinates relative to bregma: 1.4 mm AP, 0.5 mm ML (10°), and -4.5 mm DV) using a 29 G stainless steel cannula connected to a 2 μl Hamilton syringe or a Nanoject system (Drummond Scientific, PA). The injectors were retracted slowly after 5 minutes. Mice were given carprofen (5 mg/kg) post-surgery for pain relief.

### Brain slice preparation and electrophysiological recordings

After > 4 weeks of viral incubation, mice were anesthetized with Euthasol (i.p., Virbac AH, Inc, TX) and then decapitated. Brains were rapidly removed and placed in room temperature (25 °C) modified Krebs’ buffer containing (in mM): 125 NaCl, 4.5 KCl, 1.0 MgCl_2_, 1.2 CaCl_2_, 1.25 NaH_2_PO_4_, 11 D-glucose, and 23.8 NaHCO_3_, bubbled with 95/5% O_2_/CO_2_ with 5 µM MK-801 to increase slice viability. Using a vibrating microtome (Leica Biosystems, IL). Coronal brain slices (220 µm) containing the ventral striatum were collected and incubated at 32 °C for 4 min, then transferred to room temperature until use. All recordings were from neurons in the ventro-medial shell of the nucleus accumbens. Cells were visualized using IR-DIC and fluorescence on an upright Olympus BX51WI microscope (Olympus, MA). Transduced D1-MSNs were identified by visualization of eGFP or eYFP. Whole-cell patch-clamp recordings were made at 35±1 °C with a Multiclamp 700B amplifier and Digidata 1440a digitizer with Clampex and Axoscope software (Molecular Devices, CA). Data were digitized at 20 kHz. For voltage-clamp experiments, data were filtered at 2 kHz. Recordings were performed in modified Krebs’ buffer. In the majority of experiments, receptor antagonists were used to eliminate fast synaptic transmission (3 µM NBQX or DNQX and 100 µM picrotoxin). Pipette resistance was 1.5-2.5 MΩ when filled with internal solution containing (in mM) K-methylsulfate (122), HEPES (10), EGTA (0.45), NaCl (9), MgCl_2_ (1.8), CaCl_2_ (0.1), Mg-ATP (4), Na-GTP (0.3), creatine phosphate disodium (14), pH 7.35 and 284 mOsm.

Assuming permeability to Cl^-^ only, the calculated reversal potential of PSAM^4^-GlyR was -62.3 mV. Assuming that PSAM^4^-GlyR has similar Cl^-^/HCO_3_^-^ permeability to GlyR (P_HCO3-_:P_Cl-_ = 0.14) (37)), the calculated reversal potential of PSAM^4^-GlyR in our recording conditions was -60.7 mV ([Cl^-^]_out_ 133.9 mM, [Cl^-^]_in_ 12.8 mM, [HCO3^-^]_out_ 23.8 mM, and assuming [HCO3^-^]_in_ 8 mM). Series resistance was monitored throughout the recordings and not compensated. Reported voltages were corrected for a liquid junction potential of -8 mV between the internal and external solutions. To measure neuronal excitability, current was injected in 50 pA increments (2 s). In determining the current-voltage (I-V) relationship using voltage steps, the external solution of some recordings included 1 µM tetrodotoxin (TTX) to eliminate sodium channel-dependent spiking. There were no differences in the shape of the I-V with or without TTX, so data were combined. Reversal potentials were determined by linear regression considering each replicate an individual point. Recordings in which current did not cross 0 pA were omitted from analysis (*n* = 2). GABA-A receptor-mediated synaptic currents were evoked with electrical stimulation delivered by a bipolar stimulating electrode placed in the brain slice, in the presence of ionotropic glutamate receptor antagonists (NBQX/DNQX) to isolate fast GABAergic synaptic transmission. Peak amplitude of GABA-A receptor-mediated synaptic currents was measured 3-4 ms from the apparent peak to remove any potential contribution from a stimulation-induced ‘escaped’ action potential.

### Immunohistochemistry and confocal microscopy

Mice microinjected with PSAM^4^-GlyR *vs*. control AAV into the medial ventral striatum received an i.p. injection of uPSEM^792^ (3 mg/kg) or saline followed 30 minutes later by an i.p. injection of fentanyl (0.2 mg/kg) or saline (Figure 1B). After 90 min, at the expected peak of c-fos protein expression (38), mice were euthanized with Euthasol (i.p.) and transcardially perfused with 1X PBS followed by ice-cold 4% paraformaldehyde (pH 7.4, Sigma Aldrich, MO). The brains were collected and fixed overnight in 4% paraformaldehyde at 4 °C, then moved to 1XPBS. Coronal brains slices (50 μm) containing the viral injection sites were collected using a vibratome (Leica Biosystems, IL). Free-floating brain slices were washed in PBS (x3), permeabilized and blocked in a solution containing PBS, 0.3% Triton-X, and 5% normal donkey serum for 2 h. Slices were then incubated at 4° overnight in a solution containing PBS, 0.03% Triton-X, 5% normal donkey serum, and 1:4000 rabbit anti-c-fos primary antibody (Cell Signaling, MA). This was followed by a wash in PBS (x3) and incubation in 1:500 donkey anti-rabbit secondary antibody, conjugated to Alexa-Fluor-647 (Jackson ImmunoResearch, PA) for 1 h and 45 min. Slices were then washed in PBS (x3) and mounted with DAPI Fluoromount mounting medium (Thermo Fisher Scientific, MA). Confocal images were collected using an Olympus Fluoview FV1000 confocal microscope with 20x (0.75 NA) or 40x (0.95 NA) objective lens and processed using ImageJ. Z-stack images of the viral injection site in the right hemisphere (2-3 brain sections) were collected. Expression of c-fos and eGFP/eYFP was manually quantified by counting fluorescent cells within the image frame. The number of non-transduced cells was obtained by subtracting the number of transduced cells from the number of cells stained with DAPI. Results from all brain sections were averaged for each mouse.

### Drugs

uPSEM^792^, MK-801, picrotoxin, and NBQX were purchased from Tocris (MN). All other electrophysiological reagents were obtained from Sigma-Aldrich (MO).

### Data analysis

Data were analyzed using Clampfit 10.7. Data are presented as representative traces, or in bar graphs with means ± SEM, and in scatter plots where each point is an individual cell. As noted in figure legends, *n* = number of distinct cells or mice. Data were analyzed using GraphPad Prism (v 8.1.1; GraphPad software, CA). The statistical tests we used included *t*-tests (paired or unpaired), ANOVAs (with or without repeated measures) with Sidak’s or Dunnett’s multiple comparison test (as recommended by Prism), ANOVA mixed-effects analysis (when missing values prohibited repeated-measures analysis with ANOVAs), Pearson’s correlation, and linear regression (we ensured there were no departures from linearity with replicates test). Significance level was set at *p*< 0.05.

#### ACKNOWLEDGMENTS

We would like to thank Dr. Shiliang Zhang and the Confocal and Electron Microscopy Core at the NIDA Intramural Research Program for helping with confocal imaging. This project was supported by the Intramural Research Program at the National Institute on Drug Abuse, DA048085 (KM), and the Center on Compulsive Behaviors, National Institutes of Health via NIH Director’s Challenge Award (SCG).

## AUTHOR CONTRIBUTIONS

KM and SG designed the experiments. KM, MO, and AB performed the experiments. KM, MO, and SG analyzed the data and prepared the figures. KM, MO, and SG wrote the manuscript.

## REFERENCES

1. Roth BL. DREADDs for Neuroscientists. Neuron. 2016;89(4):683–94.

2. Saloman JL, Scheff NN, Snyder LM, Ross SE, Davis BM, Gold MS. Gi-DREADD Expression in Peripheral Nerves Produces Ligand-Dependent Analgesia, as well as Ligand-Independent Functional Changes in Sensory Neurons. Journal of Neuroscience. 2016;36(42):10769–81.

3. Gomez JL, Bonaventura J, Lesniak W, Mathews WB, Sysa-Shah P, Rodriguez LA, et al. Chemogenetics revealed: DREADD occupancy and activation via converted clozapine. Science. 2017;357(6350):503–7.

4. Manvich DF, Webster KA, Foster SL, Farrell MS, Ritchie JC, Porter JH, et al. The DREADD agonist clozapine N-oxide (CNO) is reverse-metabolized to clozapine and produces clozapine-like interoceptive stimulus effects in rats and mice. Sci Rep. 2018;8(1):3840.

5. Magnus CJ, Lee PH, Atasoy D, Su HH, Looger LL, Sternson SM. Chemical and genetic engineering of selective ion channel-ligand interactions. Science. 2011;333(6047):1292–6.

6. Magnus CJ, Lee PH, Bonaventura J, Zemla R, Gomez JL, Ramirez MH, et al. Ultrapotent chemogenetics for research and potential clinical applications. Science. 2019;364(6436).

7. Al-Hasani R, McCall JG, Shin G, Gomez AM, Schmitz GP, Bernardi JM, et al. Distinct Subpopulations of Nucleus Accumbens Dynorphin Neurons Drive Aversion and Reward. Neuron. 2015;87(5):1063–77.

8. Gerfen CR, Engber TM, Mahan LC, Susel Z, Chase TN, Monsma FJ, Jr., et al. D1 and D2 dopamine receptor-regulated gene expression of striatonigral and striatopallidal neurons. Science. 1990;250(4986):1429–32.

9. Krashes MJ, Shah BP, Madara JC, Olson DP, Strochlic DE, Garfield AS, et al. An excitatory paraventricular nucleus to AgRP neuron circuit that drives hunger. Nature. 2014;507(7491):238–42.

10. Chung L. A Brief Introduction to the Transduction of Neural Activity into Fos Signal. Dev Reprod. 2015;19(2):61–7.

11. Cruz FC, Javier Rubio F, Hope BT. Using c-fos to study neuronal ensembles in corticostriatal circuitry of addiction. Brain Res. 2015;1628(Pt A):157–73.

12. Sheng M, Greenberg ME. The regulation and function of c-fos and other immediate early genes in the nervous system. Neuron. 1990;4(4):477–85.

13. Hunt SP, Pini A, Evan G. Induction of c-fos-like protein in spinal cord neurons following sensory stimulation. Nature. 1987;328(6131):632–4.

14. Enoksson T, Bertran-Gonzalez J, Christie MJ. Nucleus accumbens D2- and D1-receptor expressing medium spiny neurons are selectively activated by morphine withdrawal and acute morphine, respectively. Neuropharmacology. 2012;62(8):2463–71.

15. Lobo MK, Zaman S, Damez-Werno DM, Koo JW, Bagot RC, DiNieri JA, et al. DeltaFosB induction in striatal medium spiny neuron subtypes in response to chronic pharmacological, emotional, and optogenetic stimuli. J Neurosci. 2013;33(47):18381–95.

16. Chang SL, Squinto SP, Harlan RE. Morphine activation of c-fos expression in rat brain. Biochem Biophys Res Commun. 1988;157(2):698–704.

17. Liu J, Nickolenko J, Sharp FR. Morphine induces c-fos and junB in striatum and nucleus accumbens via D1 and N-methyl-D-aspartate receptors. Proc Natl Acad Sci U S A. 1994;91(18):8537–41.

18. Planert H, Berger TK, Silberberg G. Membrane properties of striatal direct and indirect pathway neurons in mouse and rat slices and their modulation by dopamine. PLoS One. 2013;8(3):e57054.

19. Huguenard JR, Alger BE. Whole-cell voltage-clamp study of the fading of GABA-activated currents in acutely dissociated hippocampal neurons. J Neurophysiol. 1986;56(1):1–18.

20. Staley KJ, Soldo BL, Proctor WR. Ionic mechanisms of neuronal excitation by inhibitory GABAA receptors. Science. 1995;269(5226):977–81.

21. Thompson SM, Gahwiler BH. Activity-dependent disinhibition. I. Repetitive stimulation reduces IPSP driving force and conductance in the hippocampus in vitro. J Neurophysiol. 1989;61(3):501–11.

22. Karlsson U, Druzin M, Johansson S. Cl(−) concentration changes and desensitization of GABA(A) and glycine receptors. J Gen Physiol. 2011;138(6):609–26.

23. Raimondo JV, Kay L, Ellender TJ, Akerman CJ. Optogenetic silencing strategies differ in their effects on inhibitory synaptic transmission. Nat Neurosci. 2012;15(8):1102–4.

24. Alfonsa H, Merricks EM, Codadu NK, Cunningham MO, Deisseroth K, Racca C, et al. The contribution of raised intraneuronal chloride to epileptic network activity. J Neurosci. 2015;35(20):7715–26.

25. Wiegert JS, Mahn M, Prigge M, Printz Y, Yizhar O. Silencing Neurons: Tools, Applications, and Experimental Constraints. Neuron. 2017;95(3):504–29.

26. Doyon N, Vinay L, Prescott SA, De Koninck Y. Chloride Regulation: A Dynamic Equilibrium Crucial for Synaptic Inhibition. Neuron. 2016;89(6):1157–72.

27. O’Donnell P, Greene J, Pabello N, Lewis BL, Grace AA. Modulation of cell firing in the nucleus accumbens. Ann N Y Acad Sci. 1999;877(1 ADVANCING FRO):157–75.

28. O’Donnell P, Grace AA. Synaptic interactions among excitatory afferents to nucleus accumbens neurons: hippocampal gating of prefrontal cortical input. J Neurosci. 1995;15(5 Pt 1):3622–39.

29. Burke DA, Rotstein HG, Alvarez VA. Striatal Local Circuitry: A New Framework for Lateral Inhibition. Neuron. 2017;96(2):267–84.

30. Koos T, Tepper JM. Inhibitory control of neostriatal projection neurons by GABAergic interneurons. Nat Neurosci. 1999;2(5):467–72.

31. Mayer ML, Westbrook GL. Permeation and block of N-methyl-D-aspartic acid receptor channels by divalent cations in mouse cultured central neurones. J Physiol. 1987;394(1):501–27.

32. Mayer ML, Westbrook GL, Guthrie PB. Voltage-dependent block by Mg2+ of NMDA responses in spinal cord neurones. Nature. 1984;309(5965):261–3.

33. Pasek JG, Wang X, Colbran RJ. Differential CaMKII regulation by voltage-gated calcium channels in the striatum. Mol Cell Neurosci. 2015;68:234–43.

34. Clapham DE. Calcium signaling. Cell. 2007;131(6):1047–58.

35. Lisman J, Schulman H, Cline H. The molecular basis of CaMKII function in synaptic and behavioural memory. Nat Rev Neurosci. 2002;3(3):175–90.

36. Wayman GA, Lee YS, Tokumitsu H, Silva AJ, Soderling TR. Calmodulin-kinases: modulators of neuronal development and plasticity. Neuron. 2008;59(6):914–31.

37. Jun I, Cheng MH, Sim E, Jung J, Suh BL, Kim Y, et al. Pore dilatation increases the bicarbonate permeability of CFTR, ANO1 and glycine receptor anion channels. J Physiol. 2016;594(11):2929–55.

38. Barros VN, Mundim M, Galindo LT, Bittencourt S, Porcionatto M, Mello LE. The pattern of c-Fos expression and its refractory period in the brain of rats and monkeys. Front Cell Neurosci. 2015;9:72.

